# MartiniSurf: Automated Simulations of Surface-Immobilized Biomolecular Systems with Martini

**DOI:** 10.64898/2026.03.27.714767

**Authors:** Juan Carlos Jiménez-García, Fernando López-Gallego, Xabier López, David De Sancho

## Abstract

The rational design of biomolecule immobilization strategies requires molecular-level understanding of how surface properties, tethering geometry, and structural dynamics jointly influence stability and function. Recently, coarse-grained molecular dynamics simulations based on the Martini force field have emerged as an efficient framework for studying enzyme–surface interactions. However, the reproducible construction of immobilized systems with controlled orientations remains technically challenging, usually involving multiple computational tools. Here we present MartiniSurf, an open-source command-line framework for the preparation of protein and DNA systems immobilized on solid supports within the Martini paradigm. MartiniSurf integrates automated structure retrieval and cleaning, coarse graining via tools from the Martini force field software ecosystem, customizable surface generation, and biomolecule orientation based on user-defined anchoring residues, producing complete GROMACS-ready simulation systems. The framework supports both implicit restraint-based anchoring and explicit linker-mediated immobilization, including surfaces functionalized with user-defined ligands or linker-like moieties, enabling representation of mono- and multivalent attachment geometries at different modeling resolutions. Structure-based GōMartini potentials can be incorporated for proteins, while DNA systems are modeled using Martini 2. Optional substrate insertion, pre-coarse-grained complex handling, and automated solvation and ionization further extend system flexibility. By integrating these components into a unified workflow, MartiniSurf enables systematic and high-throughput *in silico* exploration of surface-tethered biomolecules and provides a robust computational platform for rational immobilization studies.

**TOC Graphic:** 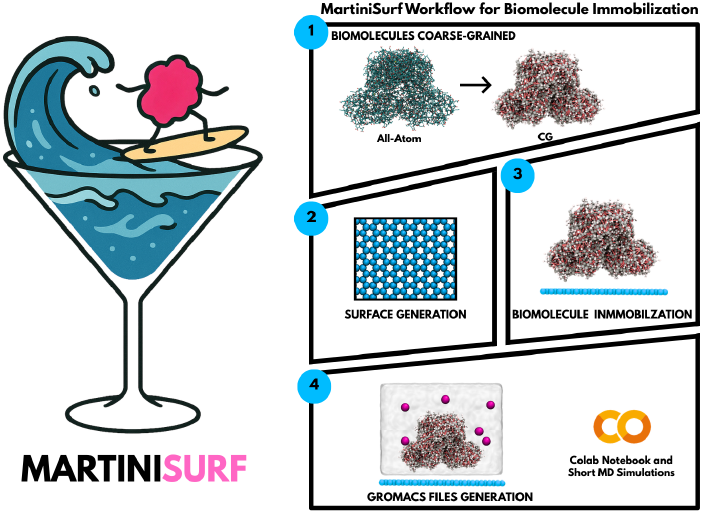

## Introduction

Despite the widespread use of biomolecule immobilization in biotechnology, developing systematic strategies to identify surface tethering sites remains a challenge. Even when identical surface chemistries are used, variations in the attachment site, orientation, or tethering geometry can lead to markedly different functional outcomes.^1–4^ As a result, there is growing interest in site-directed rational immobilization approaches that aim to control molecular orientation and local rigidity while preserving access to active regions.^5–7^ Molecular dynamics (MD) simulations offer a powerful complementary approach for probing biomolecule–surface interactions at molecular resolution. All-atom simulations, in particular, have yielded detailed insights into adsorption mechanisms, orientation preferences, and surface-induced conformational changes.^8–10^ However, their high computational cost limits systematic exploration across multiple immobilization geometries, surface models, and biomolecular variants, particularly when long timescales or large assemblies are required. Coarse-grained (CG) models address this limitation by reducing molecular resolution while retaining essential physicochemical features.^11^ Among these, the Martini force field has emerged as a widely adopted framework for large biomolecular systems.^12–16^ When combined with structure-based Gōlike potentials, Martini simulations can preserve native folds while enabling collective motions relevant to biomolecular function.^17^

Recently, we have demonstrated that structure-based Martini simulations can capture key mechanistic determinants of enzyme immobilization, including the coupling between tethering geometry, conformational dynamics, and catalytic function. ^18^ In that study, Alcohol Dehydrogenase immobilized on an agarose-like surface was simulated using the GōMartini model.^17^ Variants involving different immobilization sites and strategies were modelled through multivalent harmonic restraints on an idealized agarose-like surface, revealing orientation-dependent modulation of flexibility and substrate accessibility. However, the system construction required manual configuration steps involving manual preprocessing, custom scripts, and non-standardized orientation procedures. In particular, enforcing reproducible, orientation-controlled, and multivalent anchoring strategies — which are central to modern experimental immobilization approaches^6,19^ —remains technically challenging. This highlights the need for a unified and reproducible framework capable of systematically generating such immobilized simulation systems and extending them toward more complex surface chemistries.

To address these limitations, here we present MartiniSurf, an automated and reproducible framework for constructing surface-immobilized biomolecular systems within the Martini paradigm (Figure 1). By consolidating system preparation into a unified workflow, MartiniSurf enables systematic *in silico* exploration of immobilization geometries and provides a robust foundation for rational design studies at coarse-grained resolution.

**Figure 1:**
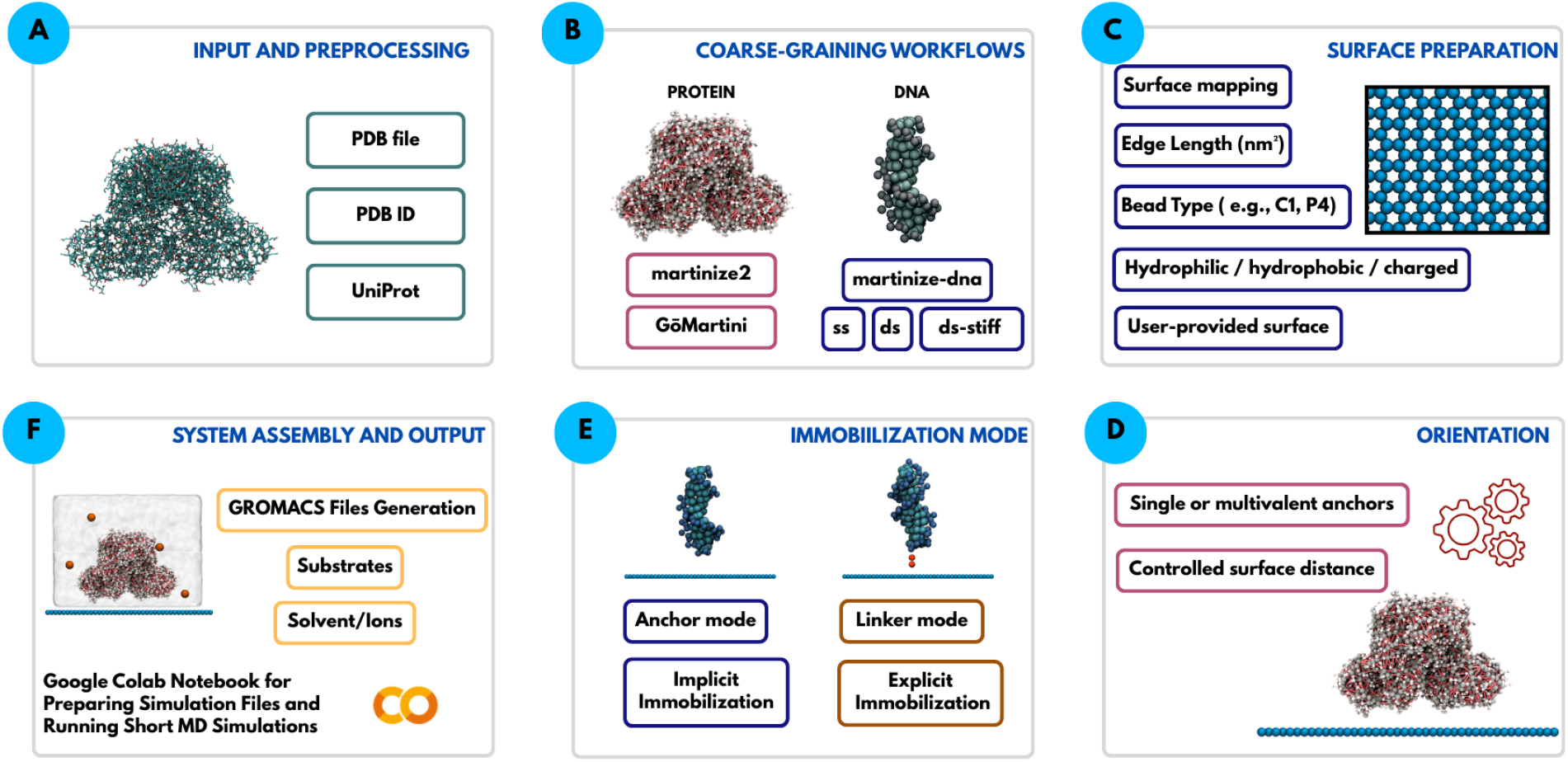
Overview of the MartiniSurf workflow for constructing surface-immobilized biomolecular systems for coarse-grained molecular dynamics. Starting from a biomolecular structure provided as a local file, PDB identifier or UniProt accession, MartiniSurf performs preprocessing, applies dedicated coarse-graining workflows for proteins or DNA, generates model surfaces with tunable mapping and bead chemistry, and orients the biomolecule relative to the interface using anchor- or linker-based immobilization strategies. The pipeline then assembles the final simulation system, including optional solvent, ions, and substrates, and produces GROMACS-ready files together with a Google Colab notebook for short molecular dynamics simulations.

## Methods and Implementation

MartiniSurf was designed around four core requirements relevant to computational studies of biomolecule immobilization. First, *automation*: the complete system-construction process should be executable from the command-line interface (ideally using a single command), minimizing manual intervention and user-dependent variability. Second, *reproducibility*: explicit input parameters define the generated system, enabling robust comparison across immobilization geometries and independent studies. Third, *flexibility*: the framework should support different types of biomolecule, force-field variants, surface models, and tethering strategies without requiring custom code modifications. Fourth, *extensibility*: the framework should be readily adaptable to incorporate new functionalities and evolving Martini model developments, ensuring long-term usability. Together, these design principles enable MartiniSurf to prepare complete simulation systems for downstream simulations using the molecular mechanics package GROMACS.^20^

### Coarse graining with tools from the Martini ecosystem

The workflow begins from a user-provided biomolecular structure specified through the `--pdb` flag, which accepts either a local `.pdb` coordinate file, a 4-character RCSB PDB identifier, or a 6-character UniProt accession. In the latter case, MartiniSurf automatically retrieves the corresponding AlphaFold structural model and converts it into the standard input required by the downstream setup pipeline.

Input structures are automatically retrieved, cleaned, and standardized prior to coarse graining. Protein systems are processed using `martinize2`,^21^ with optional elastic networks or structure-based GōMartini native-contact potentials,^17^ whereas DNA systems are handled using `martinize-dna`.^22^ At the command-line level, these alternatives are controlled through a compact set of workflow-defining flags, including `--go` for structure-based protein models, `--dna` together with `--dnatype` for nucleic-acid systems, and `--complex-config` for precoarse-grained assemblies that bypass the “martinization” stage. This design allows the same high-level interface to accommodate atomistic protein inputs, DNA structures, and pre-built coarse-grained complexes within a unified setup workflow.

### Surface generation

Following coarse graining, MartiniSurf generates model solid supports represented as Martini-bead surfaces, including built-in planar reference supports based on two-dimensional hexagonal lattices used as built-in planar reference supports. In practice, the built-in planar surface types are selected through the `--surface-mode` flag, including options such as `4-1`, `2-1`, `graphene`, and `graphite`. The lattice spacing, bead identity, charge distribution, multilayer configuration, and optional surface functionalization can be explicitly defined. In particular, MartiniSurf supports the addition of user-defined ligands or linker-like moieties to the surface through the `--surface-linkers` option, enabling the construction of chemically functionalized interfaces beyond bare reference supports. Lattice-based coarse-grained surfaces have previously been explored in Martini simulations to model carbon-based and chemically tuned materials. In these approaches, bead-type adjustments have been used to capture the effects of surface chemistry,^23^ while alternative mapping schemes (e.g., 4:1 and 2:1 representations) have been proposed for graphitic materials to capture surface density and interaction properties.^24,25^ Extensions to chemically heterogeneous graphene oxide surfaces that combine modified bead types and reduced mapping schemes have also been reported.^26^ Similar hexagonal lattice strategies have been used to represent polysaccharide-based materials, such as agarose-like surfaces, highlighting the broader applicability of this framework across distinct classes of immobilization supports.^27^ MartiniSurf formalizes these approaches, enabling controlled exploration of how surface physicochemical properties influence the dynamics of immobilized biomolecules and extending our previous mechanistic studies toward systematically tunable, chemically resolved surface models. In addition to internally generated supports, MartiniSurf also accepts user-provided surface coordinate and topology files. This flexibility allows integration of externally constructed or experimentally inspired architectures, including heterogeneous supports or previously parameterized materials.

Within the built-in surface-generation workflow, planar carbon-based materials such as graphene and graphite are available through the corresponding `--surface-mode` options, whereas carbon nanotubes are generated through dedicated CNT modes (`cnt-m2` and `cnt-m3`) in the surface-building module. MartiniSurf reuses code from earlier coarse-grained studies of carbon nanomaterials, leveraging recent Martini models for graphene, carbon nanotubes, and fullerenes to include these surfaces in the workflow, ^28,29^ thereby enabling simulations of biomolecule immobilization on chemically realistic carbon materials. Likewise, saccharide-rich supports such as agarose can be represented using dedicated Martini carbohydrate parameterizations,^30^ allowing chemically resolved descriptions of polysaccharide-based materials beyond idealized lattice models. In this way, MartiniSurf combines idealized reference surfaces with implemented carbon-based nanomaterials while remaining fully compatible with emerging Martini-based material models.

### Biomolecule orientation

A central component of MartiniSurf is its orientation engine, which aligns user-defined tethering residues toward the surface at a prescribed separation distance through rigid-body transformations. This functionality is exposed through three CLI-defined surface-coupling regimes: anchor based immobilisation, in which selected residues (–anchor) are restrained at a distance (–dist) relative to the surface; explicit linker-based immobilization (`--linker`, `--linker-group`), in which attachment is mediated by a coarse-grained linker; and adsorption mode (`--ads-mode`), which orients the biomolecule toward the surface without imposing an explicit anchoring setup. For instance, these modes can be used to define multivalent protein immobilization through repeated `--anchor` specifications (e.g., `--anchor A 8 10 11 --anchor D 8 10 11`) or DNA tethering through an explicit aliphatic linker using `--linker` together with a residue selection such as `--linker-group A 1`. Because these surface-coupling modes are fully parameter-driven, identical input configurations always generate the same coordinate and topology files, ensuring reproducible immobilization geometries.

Defining the coupling scheme before solvation and counterion addition enables precise control over orientation and supports multivalent attachment configurations consistent with experimental tethering strategies. Within this framework, anchor and linker modes provide complementary representations of immobilization. In *anchor mode*, surface attachment is modeled implicitly through harmonic restraints between selected residues and the support, reproducing the effective tethering strategy used in previous mechanistic studies of ADH immobilization.^18^ In *linker mode*, an explicit tether molecule is introduced between the surface and the biomolecule, allowing linker geometry, flexibility, and surface functionalization to be modeled directly, and enabling systematic exploration of how linker architecture and surface chemistry influence protein–surface coupling. Linker properties have been shown to play a critical role in modulating enzyme behavior at solid interfaces; for example, enzyme thermodynamics can be strongly affected by linker length and flexibility.^31^

Because linker molecules and surface-bound functional groups are treated as explicit coarse-grained components, their use requires prior Martini-compatible parametrization, including appropriate bead types, bonded terms, and interaction parameters consistent with the target force-field variant. In this context, automated small-molecule coarse-graining tools such as AutoMartini for Martini 2^32^ and recent Martini 3 workflows^33^ provide a practical route for generating linker topologies compatible with the Martini framework and can facilitate the incorporation of chemically resolved surface modifications into MartiniSurf workflows.

Representative systems generated with this workflow are shown in Figure 2. Panel (A) illustrates immobilization of the BsADH-H3 variant on an agarose-like surface via multivalent harmonic restraints (residues 8H/10H/11H), corresponding to the configuration analyzed previously.^18^ Panel (B) extends this setup toward explicitly functionalized surfaces, where surface properties become adjustable structural parameters. Panel (C) presents the complete catalytic assembly constructed with MartiniSurf, including the enzyme, the NADH cofactor,^34^ and inserted ethanol substrate molecules, mirroring the enzyme–cofactor–substrate system examined in the prior work. These examples highlight how orientation control, surface definition, and catalytic components are consistently incorporated within a unified simulation framework.

**Figure 2:**
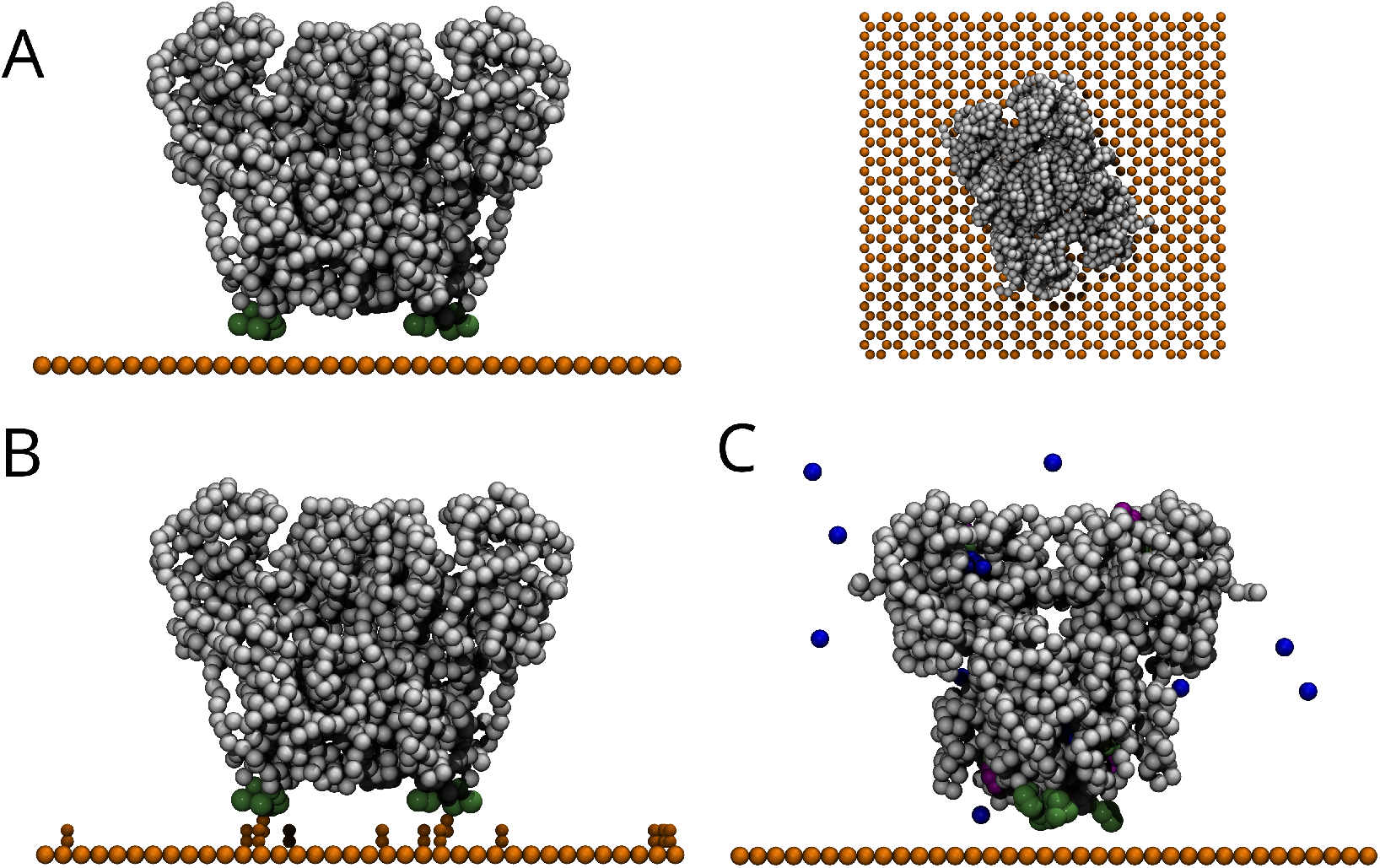
Surface-immobilized ADH systems built with MartiniSurf. (A) Immobilization on an agarose surface (side and top view). (B) Chemically functionalized surface with explicit surface-bound groups. (C) Complete catalytic system including NADH and inserted ethanol molecules. The enzyme is shown in white, the surface beads in orange, anchoring residues in green, surface functional groups in yellow, NADH in purple, and ethanol molecules in blue.

### Extension to DNA immobilization

Beyond proteins, surface-immobilized nucleic acids are also of broad interest in biotechnology, particularly in applications such as biosensing, nanobiotechnology, and DNA-based catalysis.^35,36^ In this context, aptamers —short DNA or RNA oligonucleotides capable of specifically binding molecular targets— represent an important class of surface-functional biomolecules, as their performance often depends on how tethering and surface proximity affect structure and accessibility. This makes nucleic-acid immobilization a relevant test case for extending MartiniSurf beyond enzyme–surface systems.

To assess the applicability of MartiniSurf to surface-immobilized nucleic-acid systems, we constructed a coarse-grained model reproducing the DNA–graphene setup reported by Kabeláč *et al*.^37^ In this workflow, the DNA mode is enabled through `--dna` together with the corresponding nucleic-acid model selection (`--dnatype`), while explicit tethering is introduced using `--linker` and `--linker-group`. The decamer sequence 5^*′*^-CCACTAGTGG-3^*′*^ was modeled in the Martini representation, explicitly including at the coarse-grained level the C6 aliphatic linker (–(CH_2_)_6_–) attached to the 5^*′*^ terminus. In Figure 3A, we show the two systems evaluated here, namely DNA immobilized on uncharged graphene and on positively charged graphene. As in the protein, general system-completion flags such as `--solvate`, `--ionize`, and `--salt-conc` can be used to generate fully hydrated and ionized systems for downstream GROMACS simulations. In addition, the DNA workflow supports partial conversion of solvent into frozen water through `--freeze-water-fraction` and can also be prepared with polarizable water when required by the simulation setup.

**Figure 3:**
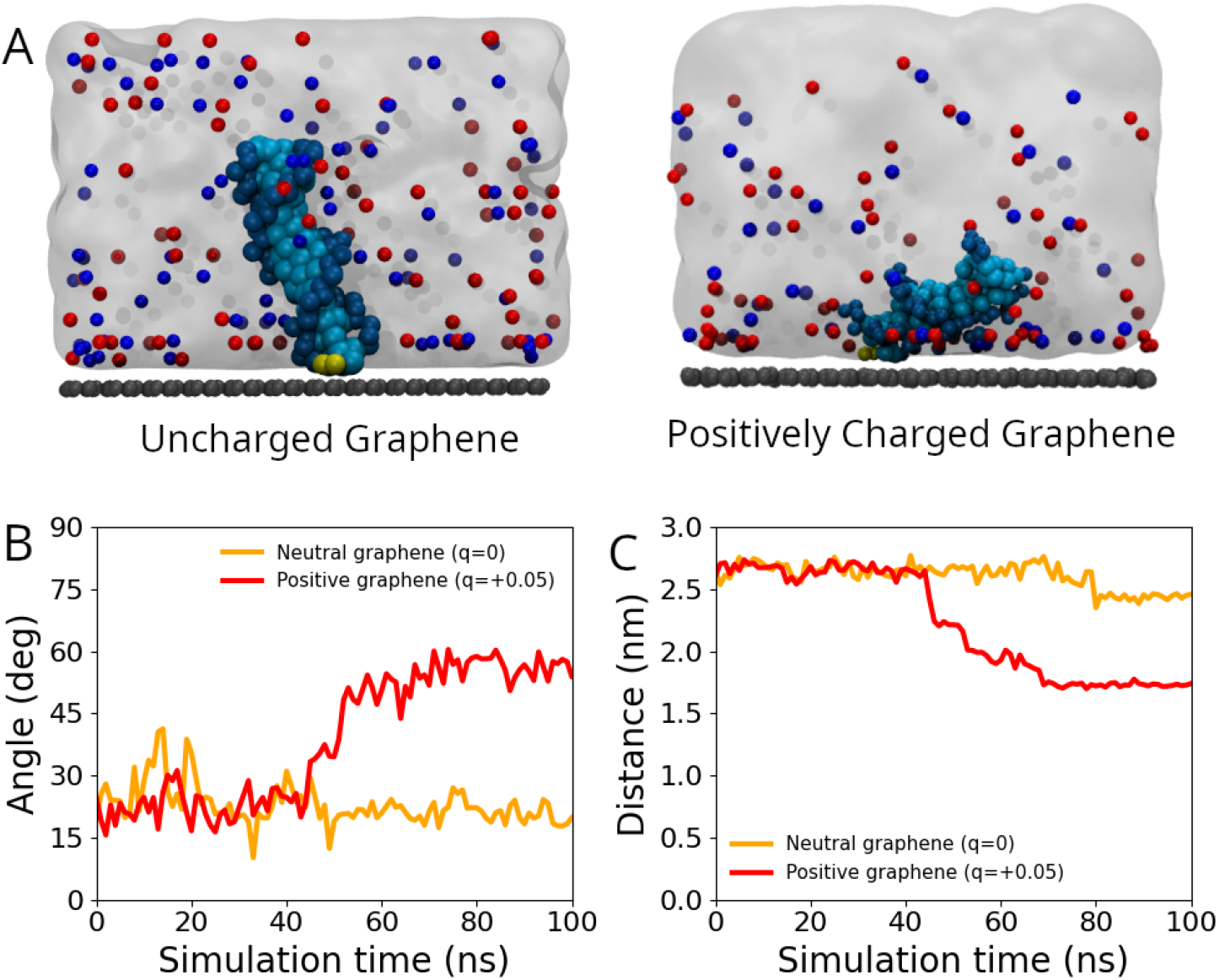
Structural stability and orientation of the surface-immobilized DNA. (A) Representative snapshot of the immobilized DNA on a graphene-like surface. The DNA is shown in a blue–cyan gradient, the surface beads in gray, and the linker in yellow. (B) Time evolution of the tilt angle between the DNA axis and the surface normal (Z-axis), where 0° corresponds to a perpendicular orientation and 90° to a parallel configuration. (C) Distance between the DNA center of mass and the graphene surface.

The coarse-grained simulations evaluated in this workflow reproduce the main orientational trends reported in the all-atom study for DNA attached to graphene surfaces of different charge. In the neutral graphene system, the DNA remains predominantly oriented away from the surface, with a relatively small tilt angle and a nearly constant center-of-mass distance over the trajectory, consistent with preservation of the anchored geometry. In contrast, on positively charged graphene, the DNA progressively bends toward the surface, showing a marked increase in tilt angle together with a reduction in the DNA–surface distance (Figure 3B,C). This behavior is consistent with the atomistic description that uncharged graphene favors a more upright configuration, whereas positive surface charge promotes closer DNA–surface association through electrostatic attraction. Overall, these results indicate that MartiniSurf captures the essential charge-dependent positional and orientational response previously described at all-atom resolution, while enabling efficient simulations at extended temporal and spatial scales. ^37^

### Command-line syntax and output

MartiniSurf is invoked through a unified command-line interface in which the user specifies the biomolecular input, surface-construction parameters, orientation mode, and optional post-processing steps through dedicated flags. For clarity, the main MartiniSurf commandline flags can be grouped according to their functional role in the setup workflow, as summarized in Table 1.

**Table 1:** Representative MartiniSurf flags by function.

| Function | Main flags |
| --- | --- |
| Input structure | --pdb, --complex-config |
| Protein setup | --moltype, --go, --dssp |
| DNA setup | --dna, --dnatype |
| Surface definition and functionalization | --surface-mode, --surface-bead, --dx, --lx, --ly, --surface-layers, --surface-linkers |
| Anchor-based immobilization | --anchor, --dist |
| Linker-based immobilization | --linker, --linker-group |
| Adsorption setup | --ads-mode |
| System completion | --solvate, --ionize, --salt-conc |
| DNA-Martini 2 Water | --freeze-water-fraction, --freeze-water-seed |

The final output of MartiniSurf is a fully assembled, GROMACS-ready simulation directory containing coordinate files, topology definitions, index groups, anchoring or linker restraint files when applicable, and template molecular dynamics parameter files. The default output structure is organized into topology, MDP, and system folders, allowing immediate integration into downstream equilibration and production workflows.

### Representative command-line examples

Representative MartiniSurf workflows can be specified using a small number of high-level flags. For example, anchor-based protein immobilization with solvation and ionization can be generated with:

`martinisurf --pdb inputs/1RJW.pdb --dssp --go --moltype Protein \`

`--surface-mode 4-1 --surface-bead P4 --dx 0.47 --lx 15 --ly 15 \`

`--anchor A 8 10 11 --anchor D 8 10 11 --dist 1.0 \`

`--solvate --ionize --salt-conc 0.15 --merge A,B,C,D`

A DNA linker-based workflow including solvation, ionization, and frozen water can be generated with:

`martinisurf --dna --dnatype ds-stiff --pdb inputs/4C64.pdb \`

`--surface-mode 4-1 --lx 10 --ly 10 --dx 0.27 \`

`--surface-layers 2\ --surface-bead C1 C1\`

`--linker inputs/ALK.gro --linker-group A 1 \`

`--solvate --ionize --salt-conc 0.15 \`

`--freeze-water-fraction 0.10 --freeze-water-seed 42 --merge A,B`

## Applications and Outlook

MartiniSurf enables systematic extension of previously validated immobilization models toward chemically resolved and mechanistically richer surface representations. By lowering the technical barrier to constructing surface-immobilized coarse-grained systems, the framework allows controlled variation of anchoring geometry, surface lattice properties, and linker architecture within a fully reproducible workflow. This design facilitates quantitative investigation of how surface physicochemical characteristics modulate conformational stability, dynamic coupling, and catalytic accessibility in immobilized enzymes.

While MartiniSurf includes planar reference surfaces for controlled mechanistic studies, the current implementation also supports graphitic materials such as graphene, graphite, and carbon nanotubes. This combination enables both systematic comparisons on simplified supports and simulations on chemically relevant carbon-based materials within the same framework. In addition, the modular design of MartiniSurf enables straightforward incorporation of alternative surface architectures, heterogeneous materials, and externally parameterized geometries. In principle, similar extensions could also be considered for softer lipid-based interfaces, given the broad use of Martini in lipid bilayer simulations.^12,13,38^

Future developments may incorporate additional surface chemistries, dynamically flexible supports, optimized contact-map definitions for GōMartini models,^39^ alternative structure-preserving schemes such as OLIVES for Martini proteins,^40^ and automated post-processing and analysis pipelines, further strengthening MartiniSurf as an extensible platform for exploring surface-immobilized biomolecular systems.

## Implementation

MartiniSurf is implemented in Python and distributed as an open-source package. The codebase follows a modular architecture in which structure processing and coarse graining, surface construction, orientation, and final system assembly are handled by separate components coordinated through a high-level command-line interface. This separation of responsibilities improves maintainability, facilitates extension to new surface geometries and tethering strategies, and supports integration into automated and high-throughput simulation pipelines.

MartiniSurf interfaces with established external programs, including `martinize2` for proteins^21^ or `martinize-dna` for nucleic acids,^22^ DSSP,^41^ MDtraj,^42^ and GROMACS.^20^ Because all outputs are generated from explicit input parameters, MartiniSurf promotes reproducible research and transparent methodological workflows.

To improve accessibility and interoperability within the Martini ecosystem, MartiniSurf also provides cloud-based Google Colab notebooks for protein and DNA system preparation. These interactive environments incorporate automated workflows for Martini 2^32^ and Martini 3,^33^ enabling atomistic-to-coarse-grained mapping, bead-type assignment, and topology generation for small molecules without requiring local software installation. In addition to system building, the notebooks support short molecular-dynamics simulations directly in Colab, facilitating rapid structural validation and consistency checks before large-scale production runs. This functionality complements the command-line interface and broadens the accessibility of the package across computational environments.

The source code, documentation, and usage examples are available at https://github.com/BioKT/MartiniSurf and https://biokt.github.io/MartiniSurf/.

## Acknowledgement

The work has been financed by grants METACELL 818089 and FET-OPEN (HOTZYMES 829162) from ERC-Co, grant PID2024-158678NB-I00 funded by MICIU/AEI/10.13039/501100011033 and “ERDF A way of making Europe”. The authors acknowledge financial support received from the IKUR Strategy under the collaboration agreement between Ikerbasque Foundation and DIPC on behalf of the Department of Science, Universities and Innovation of the Basque Government. The authors also thank the IZO-SGI SGIker (UPV/EHU/ERDF,EU) DIPC for technical and human support and for the allocation of computational resources.

